# Antigenic variation impacts gonococcal lifestyle and antibiotic tolerance by modulating interbacterial forces

**DOI:** 10.1101/2023.07.06.548055

**Authors:** Isabelle Wielert, Sebastian Kraus-Römer, Thorsten E. Volkmann, Lisa Craig, Paul G. Higgins, Berenike Maier

## Abstract

Type 4 pili (T4P) are multifunctional filaments involved in adhesion, surface motility, colony formation, and horizontal gene transfer. These extracellular polymers are surface-exposed and, therefore, act as antigens. The human pathogen *Neisseria gonorrhoeae* uses pilin antigenic variation to escape immune surveillance, yet it is unclear how antigenic variation impacts other functions of T4P. Here, we addressed this question by replacing the major pilin of a laboratory strain of *N. gonorrhoeae* with pilins from clinical isolates. Structural predictions reveal filament features that vary from one strain to the next, with the potential to impact pilus:pilus interactions. Using a combination of laser tweezers, electron microscopy, and advanced image analysis, we explore the phenotypic consequences of these structural changes. We reveal that strains differing only in their major pilin sequence vary substantially in their attractive forces, which we attribute to variations in the stereochemistry of the T4P filament. In liquid culture, strongly interacting bacteria form colonies while weakly interacting bacteria retain a planktonic lifestyle. We show that lifestyle strongly affects growth kinetics and antibiotic tolerance. In the absence of external stresses, planktonic bacteria grow faster than colony-forming bacteria. In the presence of the antibiotics ceftriaxone and ciprofloxacin, the killing kinetics indicate strongly increased tolerance of colony-forming strains. We propose that pilin antigenic variation produces a mixed population containing variants optimized for growth, colonization, or survivability under external stress. Different environments select different variants, ensuring the survival and reproduction of the population as a whole.

**Significance statement:** *Neisseria* are highly successful human pathogens that continuously vary their surface structures to escape immune surveillance. Antigenic variation of the major pilin subunit causes variations of the structure of the Type 4 pilus, a surface exposed virulence factor. Here, we investigate the effect of pilin antigenic variation on bacterial lifestyle and tolerance against antibiotics. We find that pilin antigenic variation causes changes in the physical interactions between the bacteria, resulting in distinct aggregating and planktonic phenotypes. During treatment with antibiotics, aggregating strains are more tolerant than planktonic strains by an order of magnitude. Since tolerance tends to facilitate resistance development, pilin antigenic variation reduces the efficiency of antibiotic treatment.

## Introduction

Type 4 pili (T4P) are filamentous cell appendages generated by a variety of pathogenic bacteria including the *Neisseria* species, *Pseudomonas aeruginosa*, *Vibrio cholerae*, and *Acinetobacter baumannii*. They support functions such as adhesion, motility, aggregation, and horizontal gene transfer, which are crucial for survivability, colonization, and virulence [1,2]. In pathogenic *Neisseria* the amino acid sequence of the T4P major pilin, PilE, continually changes as a result of antigenic variation [3,4]. This variability is important for escape from immune surveillance [5,6] but its roles in other cellular processes are poorly characterized.

T4P are polymers of the major pilin. They protrude from a complex that spans the inner and outer membrane [2]. Driven by cytosolic ATPases, the pilus filament elongates by polymerization and retracts by depolymerization. The filament is polymerized from major pilin subunits, with multiple low-abundance minor pilins [2]. In *N. gonorrhoeae*, the gene encoding the major pilin PilE is hypermutable by antigenic variation. Pilin antigenic variation was originally discovered by the observation that the morphology of gonococcal colonies on agar plates is related to pilus density [3,5,6]. The unpiliated phenotype was mainly caused by an antigenic variation that generated a truncated PilE, e.g. by a frameshift mutation [7,8]. In general, the macroscopic phenotype, namely flat versus spherical colony morphology, can be linked to the strength of interactions generated by the T4P of neighboring cells [9]. Antigenic variation occurs at a rate of 1.7 · 10^−3^events/cell/generation for the gonococcal lab strain MS11 used in this study [4,3,10,11]. Thus, the probability of different variants being present during a gonococcal infection is high. Segments of one of the up to 18 different *pilS* sequences [12,13] recombine with *pilE* and replace extended stretches of the *pilE* sequence. The variability along the sequence is heterogeneous and the *pilE* sequence can be subdivided into conserved, semi-variable and hyper-variable regions [14]. Pilin antigenic variation relies on a G4 motif upstream of *pilE*; deletion of this motif abolishes antigenic variation [15]. Using a mutational screen of the 3’ region of the gonococcal *pilE* coding sequence, it was shown that various point mutations caused loss of piliation [8]. For closely related *N. meningitidis* T4P, mutational analysis of *pilE* was used to generate a functional map of the major pilin [16]. In that study, a specific major pilin was used as a reference structure and single amino acid changes in different regions were associated with different T4P functions including biogenesis, adhesion, and aggregation [16]. However, it remains elusive how antigenic variations, which occur naturally and extend along dozens of amino acid residues throughout the PilE sequence, affect T4P functionality [4].

In this study, we focus on the following T4P-related functions: twitching motility, generation of attractive forces between cells, colony formation, and antibiotic tolerance. Twitching motility is a mode of surface motility driven by cycles of T4P elongation, surface attachment, and retraction [17,2,18]. *N. gonorrhoeae* follows a tug-of-war mechanism during twitching motility [19,20] and, as a consequence, gonococcal motility can be described as a correlated random walk. While T4P surface attachment mediates motility, pilus:pilus binding between adjacent cells cause attraction between gonococci [21]. In solution, this attraction causes rapid aggregation of *Neisseria* species into spherical colonies comprising thousands of cells [21–23]. The strength of the attractive force determines whether the colonies behave as liquids or solids [21,24,25], which impacts tolerance to antibiotics [26]. Attractive forces are very sensitive to the posttranslational surface modification of the T4P [24], but so far it is unclear whether the PilE and pilus structure also affect the attractive forces [27,28]. Here, we investigate how different variants of PilE impact the gonococcal lifestyle.

The emergence of antimicrobial resistance to several classes of antibiotics has made *N. gonorrhoeae* a multidrug-resistant pathogen [29]. While the mechanisms conferring gonococcal drug resistance are fairly well understood [30,31], very little is known about antibiotic tolerance of this pathogen. Tolerance describes the ability of bacteria to survive antibiotic treatment for extended periods of time [32]. This extended survival time is problematic for eradication of the pathogen and, furthermore, often precedes antibiotic resistance [33]. The mechanisms influencing tolerance are multifaceted including reduced permeability to antibiotics, reduction of growth rate and metabolic activity, membrane polarization, as well as the activation of stress responses [34]. A major tolerance mechanism is aggregation and biofilm formation [35]. Gonococcal aggregation enhances tolerance against ceftriaxone, a β-lactam targeting cell wall synthesis [31], with the physical properties of the colony impacting the degree of tolerance [26]. Ceftriaxone is currently recommended for gonorrhea treament. Since pilin antigenic variation potentially affects gonococcal aggregation and the physical properties of the colonies, it likely impacts antibiotic tolerance.

In this study, we investigate how variation of the pilin sequence affects gonococcal properties, behavior and lifestyle. Based on the PilE sequences, we predict structural changes to the surface of the T4P filament that likely impact the aggregative behavior. Using laser tweezers, we confirm that pilin variation strongly affects the attractive force between pairs of cells. We show that all *pilE* mutants in this study produce functional T4P but cluster into two distinguishable phenotypic lifestyles, planktonic and aggregating bacteria, depending on the T4P-mediated attractive force. We show that growth and survival under antibiotic treatment strongly depend on the pilin variant. Taken together, we reveal that pilin antigenic variation generates aggregative and non-aggregative T4P variants with phenotypic consequences that optimize bacterial proliferation and survival under a variety of conditions including antibiotic treatment.

## Results

### Antigenic variation alters the stereochemistry of key interaction sites on T4P

We sought to understand how pilin antigenic variation affects the physical interactions between bacteria and to characterize the resulting phenotypic changes including growth, antibiotic resistance, and antibiotic tolerance. In the first step, we cloned the *pilE* genes from the *N. gonorrhoeae* clinical isolates Ng24, Ng17 and Ng32 into the background strain, wt*\**, in place of the native *pilE* gene. wt* is *N. gonorrhoeae* MS11 with the G4 motif deleted to prevent further antigenic variation of *pilE* [64][15]. To predict whether the attractive forces and aggregative tendencies differ among the *pilE* variants, we examined their pilin sequences and predicted T4P structures. The amino acid sequences of the pilin variants were compared to that of the wt strain (Fig. 1A). Secondary structure and features were assigned based on alignment with the *N. gonorrhoeae* C30 PilE crystal structure [36] which is 90% identical in sequence to wt PilE and 99% identical in the first 120 amino acids. The sequence of the N-terminal α-helix, α1, is identical for all pilins, with the exception of a glycine instead of glutamate at position 49 of Ng17 PilE (PilE_17_). PilE_wt_ and PilE_24_ share a serine at position 63, which is post-translationally modified with a glycan in *N. gonorrhoeae* MS11 and C30 [36]. PilE_32_ and PilE_17_ have charged residues at this position. Sequence variation is observed in the αβ-loop, which lies between α1 and strand β1, in β1, in the β1-β2 loop, in the hypervariable β-hairpin, and in the C-terminal tail. We determined the overall pilin sequence identities of the variant to that of MS11 (Table S1). PilE_24_ is 90 % identical to PilE_wt_, whereas PilE_32_ is 86.6 % identical and PilE_17_ is 80.6% identical.

**Fig. 1.**
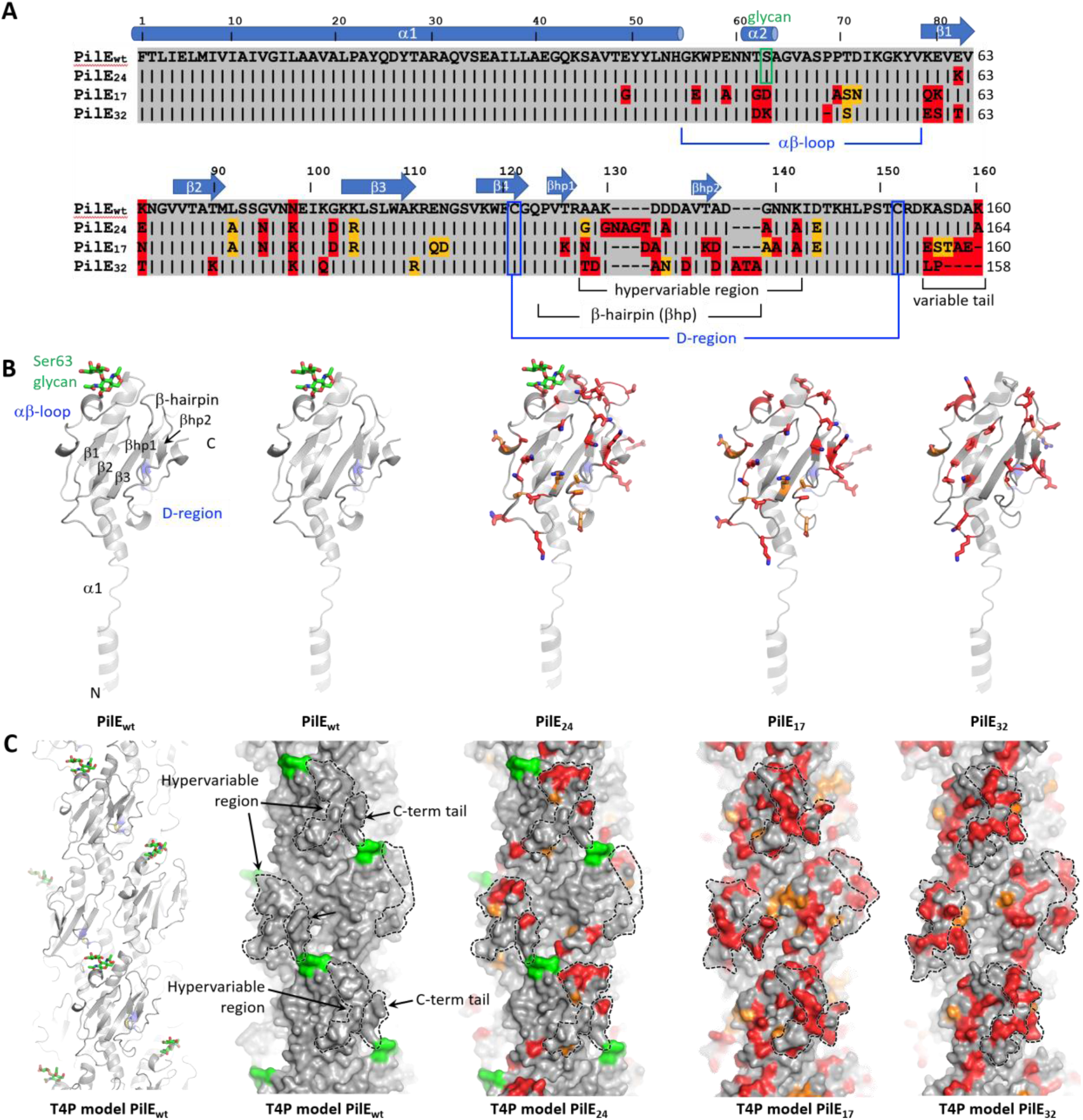
Sequence alignment and structure predictions for PilE variants and pilus filaments. A) PilE amino acid sequence alignment for wt and variant strains. Amino acids of the PilE variants that are identical to those in PilE_wt_ are shown as black bars with grey shading, conservative amino acid changes are indicated with orange shading and non-conserved residues have red shading. The secondary structure and other hallmark features of *Neisseria* Type 4 pilins are indicated based on alignment of PilE_wt_ with C30 PilE (Protein Data Bank ID 2HI2). B) Pilin models were generated using AlphaFold2 [39] (PMID: 34265844). Residues that differ from those in PilE_wt_ are shown in stick representation, with carbons colored as in Fig. 1A, nitrogens in blue and oxygens in red. Glycan carbons are green. The conserved disulfide-bonded cysteines that delineate the D-region are blue. Variable residues are located on the face of the C-terminal globular domain. C) Filament models were generated by superimposing the pilin predictions on the *N. gonorrhoeae* T4P structure (EMD-8739) and applying its symmetry parameters. The wt T4P model is shown on the left in cartoon representation and all models are shown in surface representation, colored as in Fig. 1B. The hypervariable β-hairpin and C-terminal tail together form a protruding “knob” on each subunit (dashed lines), and deep cavities or “holes” lie at the interface between subunits.

Pilin models were generated for each variant using AlphaFold2 [36]. The continuous N-terminal α-helix, α1, seen in the crystal structure of PilE [36] was replaced with the partially melted α-helix seen in the cryo-electron microscopy reconstruction of the intact pilus [37][38] (PMID: 28877506) (Fig. 1B). All variable residues are located on the face of the PilE globular domain that is exposed in the pilus filament, as expected for antigenic variation. PilE_17_ shows the greatest degree of variability, particularly in the protruding β-hairpin and C-terminal tail.

Pilus filament models were generated by superimposing each pilin model onto a single subunit in the cryoEM structure of the *N. gonorrhoeae* T4P and applying its symmetry parameters (Fig. 1C). Filament models are shown in Fig. 1C. The T4P surface is undulating: the β-hairpin and the C-terminal tail, regions, which show the highest sequence variability, together form protruding “knobs”; and deep cavities or grooves lie between the globular domains, lined with the variable features of the C-terminal tail on one side and the αβ-loop and β1-β2 loop on the other. Accordingly, the conformation of both the knobs and the cavities/grooves (“holes”) vary substantially among the variants, with wt and Ng24 T4P being most similar. The electrostatic surface potential also differs substantially from one pilus variant to the next (Fig. S1), with positively charged residues framing the holes of PilE_wt_ and PilE_24_ and negatively charged residues lining the edges of the PilE_17_ and PilE_32_ holes. The knobs of each pilin differ in their distribution of charged residues, with PilE_32_ having the most negatively charged knob. Pilus:pilus interactions likely require both structural and chemical complementarity between these surface features to allow the knobs in one filament to fit into the holes in another. Thus, these marked differences in stereochemistry among the pilus variants would impact pilus:pilus interactions and bacterial aggregation.

### Replacing the native pilin in the wt strain with antigenic variants affects bacterial attractive forces and colony formation

We examined the effects of the variations in pilus surface stereochemistry on pilus:pilus interaction and aggregation. Using a dual laser tweezers assay, we investigated the attractive forces generated by the pilin variants. We trapped pairs of bacteria for each strain and measured the T4P-mediated interaction forces between them (Fig. 2A). When bacteria in different traps interact via their T4P, and at least one T4P retracts, the cell bodies approach each other. Bacteria are deflected by a distance *d* from the centers of the traps. As the deflection increases, the restoring force of the laser traps increases as well leading to a rupture event as the optical force exceeds the (rupture) force that the pilus:pilus bond can sustain.

**Fig. 2.**
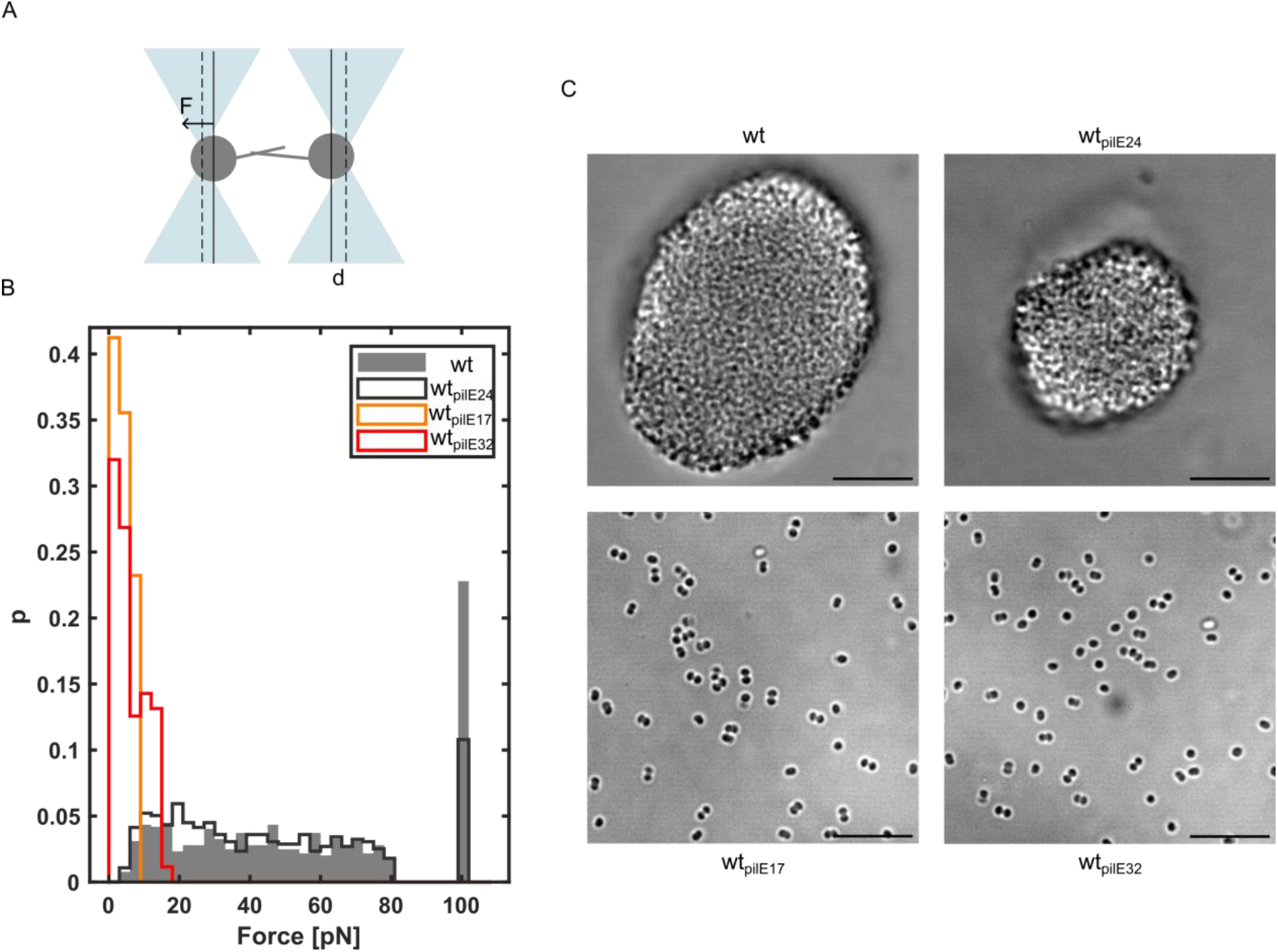
Attractive force generated between pairs of bacteria depends on the pilin sequence. A) Sketch of dual laser tweezers setup (2.64 µm distance, stiffness: *k_100%_* = 0.1 pN/nm). B) Distribution of rupture forces (number of interacting cell pairs: N_wt_ = 54, N_wt*pilE24*_ = 59, N_wt*pilE17*_ = 30, N_wt*pilE32*_ = 54). The linearity of the laser trap is limited to 80 pN. All rupture events exceeding this force were grouped into a single bin shown at 100 pN. C) Typical brightfield images of gonococci in liquid culture. Scale bar: 10 µm.

The force at which a pilus:pilus bond ruptures was used as a measure for the attractive force between gonococci. *F_rupture_* is defined as the maximal force attained before the bond breaks and the bacteria move back to their equilibrium positions. We did not observe significant differences between wt* and wt*_pilE24_* (Kolmogorow-Smirnow test (KS-test)), where the mean rupture forces were 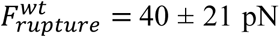 and 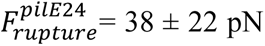, respectively (Fig. 2B). The deviation from previously published results for the wt* [26] most likely results from a different distance of the traps. Interestingly, the wt*_pilE17_* and wt*_pilE32_* variants mediate much lower rupture forces of 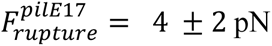 and 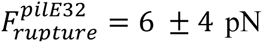, respectively. Moreover, the probability that pilus:pilus bonds form is lower for all *pilE* variant strains compared to wt* (Fig. S2).

The attractive force generated between bacteria initiates aggregation into colonies. We investigated whether the different *pilE* variants support colony formation in solution. In agreement with the dual laser tweezers experiments, we found that only the strains that mediate the strongest attractive forces, wt* and wt*_pilE24_*, are capable of in-solution colony formation (Fig. 2C). Using confocal microscopy, we investigated the effects of pilin variation on local structure and fluidity of the colonies formed by wt* and wt*_pilE24_* (Fig. S3). We found that the local order of bacteria residing in colonies was fluid-like with comparable nearest neighbor distances (Fig. S3C). Furthermore, we investigated the within-colony motility via single-cell tracking, as described previously [26]. The motility is higher inside the colony for wt*_pilE24_* than for the wt* strain, correlating with fewer interactions of single bacterial pairs (Fig. S3D). Taken together, these results show that antigenic variation of the T4P alters the attractive force between pairs of cells. Strongly interacting strains form colonies while weakly interacting strains remain planktonic.

### PilE variant strains show comparable levels of piliation and enhanced twitching motility

We hypothesize that the attractive forces between the variant strains differ because the surfaces of the T4P filaments are different (Fig. 1). Alternatively, the T4P density or T4P dynamics may cause differences in gonococcal aggregation. To distinguish between these possibilities, we first assessed the density of T4P for each strain using negative-stain transmission electron (TEM) microscopy (Fig. 3A, Fig. S4). Qualitative and quantitative comparison of the images shows that, although strain wt*_pilE32_* had a slightly higher T4P density, neither length of the pili nor their quantities differ substantially due to pilin variations. Most importantly, the planktonic strains do not show lower levels of piliation than the colony-forming strains, indicting that the piliation pattern does not cause the different aggregation phenotypes.

**Fig. 3.**
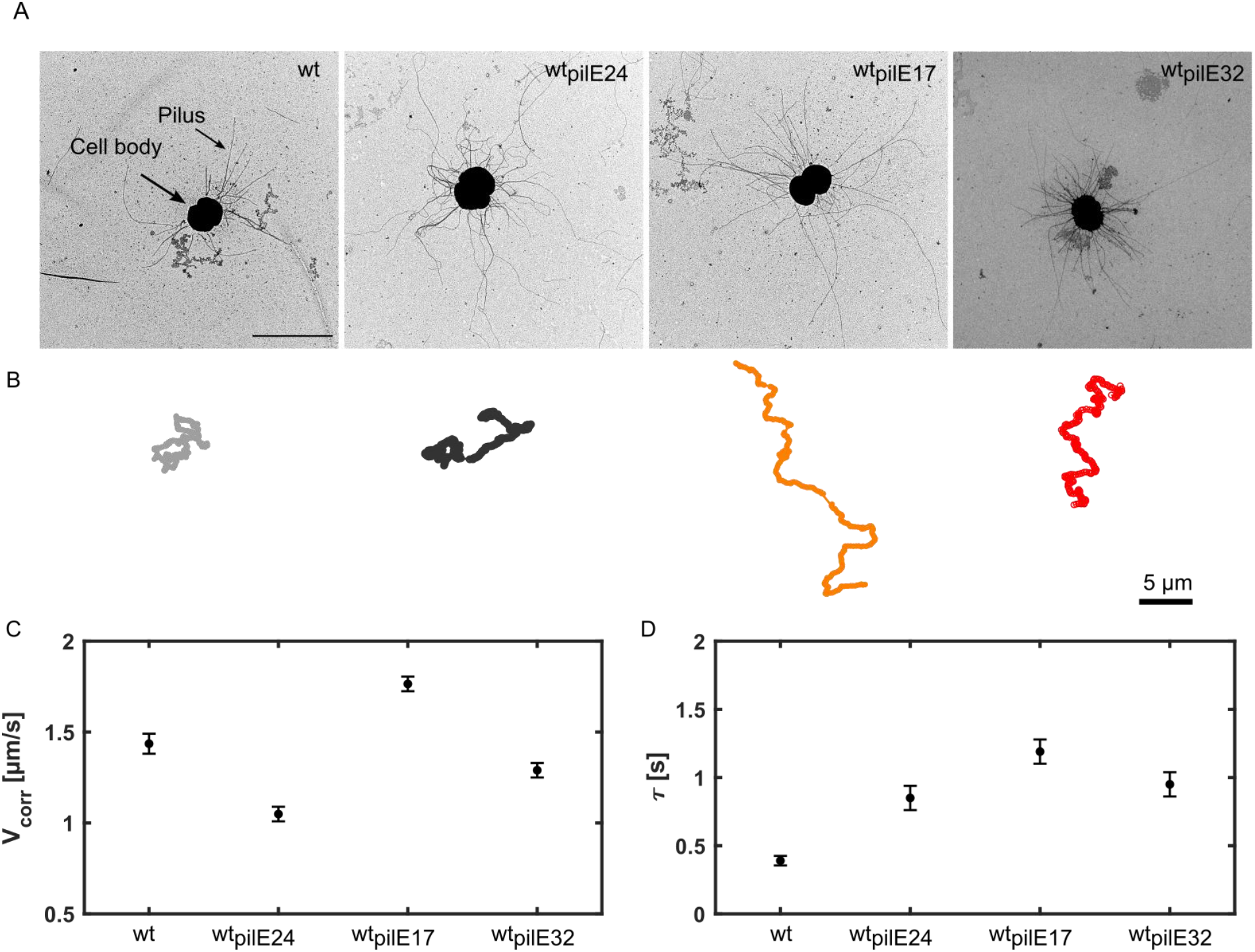
Replacing the native *pilE* sequence by antigenic variants does not reduce T4P density and maintains twitching motility. A) Representative TEM images of pilin variants. Scale bar: 2 µm. B) Trajectories of motile bacteria. Representative trajectories of pilin variants over 30 seconds (tracks made via Trackmate Image J [40]) (grey: wt*\** (Ng150), dark grey: wt*_pilE24_* (Ng242), orange: wt*_pilE17_* (Ng240), red: wt*_pilE32_* (Ng230)). Scale bar: 5 µm. C) Velocity of twitching motility for pilin variants. D) Correlation time of motile cells on a BSA coated cover slide. Error bars: 95% confidence bounds from fit of correlated random walk model.

We next compared T4P dynamics among the variants. T4P dynamics correlates quantitatively with twitching motility; the faster T4P retract the faster gonococci move on BSA-coated glass [19,20]. To find out whether variations in the PilE sequence affect T4P dynamics, we compared the trajectories of gonococci moving at the glass surface. All variants exhibit twitching motility on BSA-coated glass surfaces. A representative track for each *pilE* variant is shown in Fig. 3C. A detailed analysis of the trajectories using a correlated random walk model [20] (described in Methods, Fig. S5) showed that strain wt*_pilE17_* is the most motile; both the velocity v_corr_ and the correlation time τ were significantly higher than for the other strains (Fig. 3D, E). Moreover, the velocity shows variance among the strains with *pilE* variations (Fig. 3C), indicating that the motility characteristics depend on the specific *pilE* sequence. We conclude that all PilE variants have comparable levels of pili and generate dynamic T4P. Therefore, we attribute the effects of PilE sequence to changes in the surface of the T4P filament.

### Pilin antigenic variation affects growth and antibiotic tolerance

We tested whether the two distinct phenotypes, aggregating and planktonic gonococci, governed by pilin antigenic variation correlate with bacterial growth as well as tolerance and resistance under antibiotic treatment.

First, we characterized the growth kinetics of all *pilE* variant strains. The optical density (OD) of growing cultures is often used to characterize growth rates. We determined the OD_600_ curves in liquid culture and found complex shapes (Fig. 4A). Below an of OD_600_ 0.1, changes in cell density are undetectable due to the resolution of the instrument. Beyond this OD_600_, the planktonic strains wt*_pilE17_* and wt*_pilE32_* exhibit exponential growth up to ∼ 10 h then transition to stationary growth. In contrast, the colony forming strains wt* and wt*_pilE24_* show a very slow rise of OD_600_ for 12 h and then exhibit a sudden near-exponential increase. We interpret this to mean that wt* and wt*_pilE24_* initially grow in colonies that settle to the surface and do not contribute to the OD_600_. Around 12 h, they are released to grow planktonically, thereby increasing the OD_600_. A rapid increase in optical density is a good indicator for dissociation of *Neisseria* from colonies [49]. To test our hypothesis that wt* and wt*_pilE24_* initially grow in colonies, we determined the colony forming units (CFUs, Fig. 4B). Based on these measurements, both the planktonic and the colony forming strains exhibit exponential growth up to ∼ 10 h, after which all strains entered stationary phase. Nonetheless, the carrying capacities, i.e. the cellular concentrations, during this stationary phase differ significantly between colony-forming and planktonic strains (Fig. S6). Additionally, we determined the growth rates for all *pilE* variant strains via a linear fit to log-plotted data (Fig. 4B, C). The biofilm-forming strains have growth rates of *r*^*wt*^ = 0.57 ± 0.03 h^-1^ and *r*^*pilE*24^ = 0.46 ± 0.11 h^-1^ while the growth rates of the planktonic strains are about 25% faster, *r*^*pilE*17^ = 0.71 ± 0.12 h^-1^ and *r*^*pilE*32^ = 0.72 ± 0.10 h^-1^. After 12 h growth, the CFU values of the colony-forming strains increased exponentially and eventually reached the carrying capacity of the planktonic strains. This behaviour agrees with the optical density measurements and can be explained by dissociation of the colonies.

**Fig. 4.**
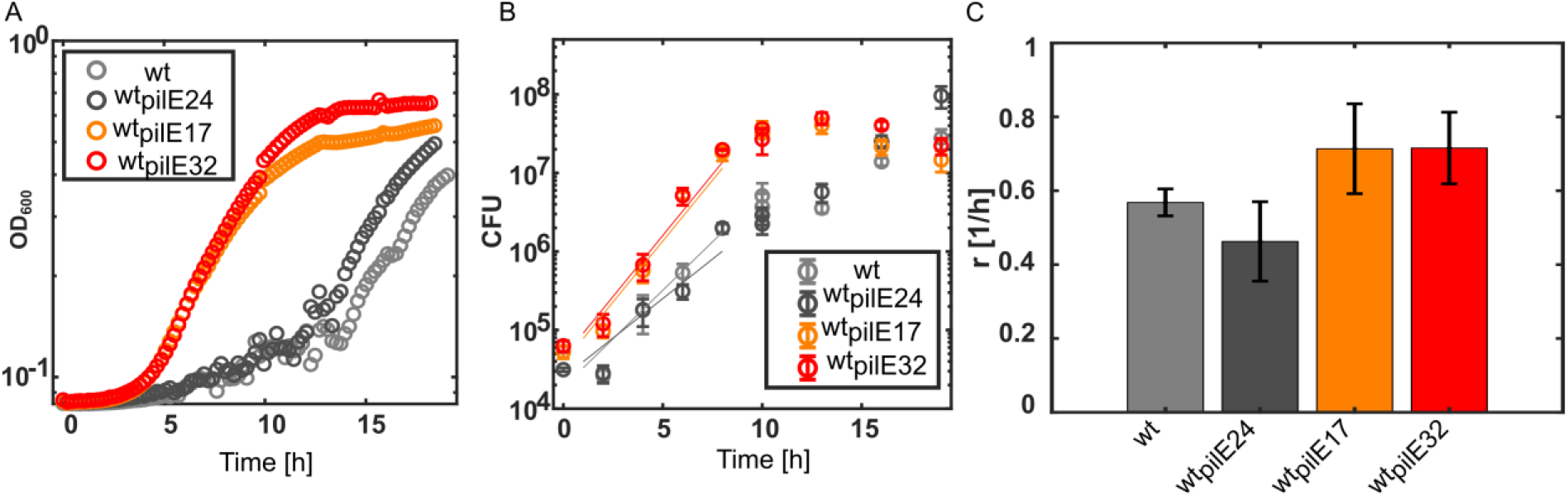
Growth kinetics of planktonic and colony forming strains. (A) Optical densities OD_600_ of *pilE* variant strains. One representative measurement out of three independent experiments is shown. (B) Colony forming units (CFU) of *pilE* variant strains. Full lines: linear fit to log-plotted data. N = 3. Error bars: standard error. C) Growth rates r determined from the linear fit shown in (B). Error bars: errors of the fits.

Next, we assessed whether variation of *pilE* affects antibiotic resistance by determining the minimal inhibitory concentrations (MICs) of antibiotics with different targets, in particular cell wall synthesis (ceftriaxone), DNA gyrase / topoisomerase (ciprofloxacin), and the ribosome (kanamycin) (Fig. 5A-C). Despite the differences in colony formation the MICs differ only slightly between the variant strains.

**Fig. 5.**
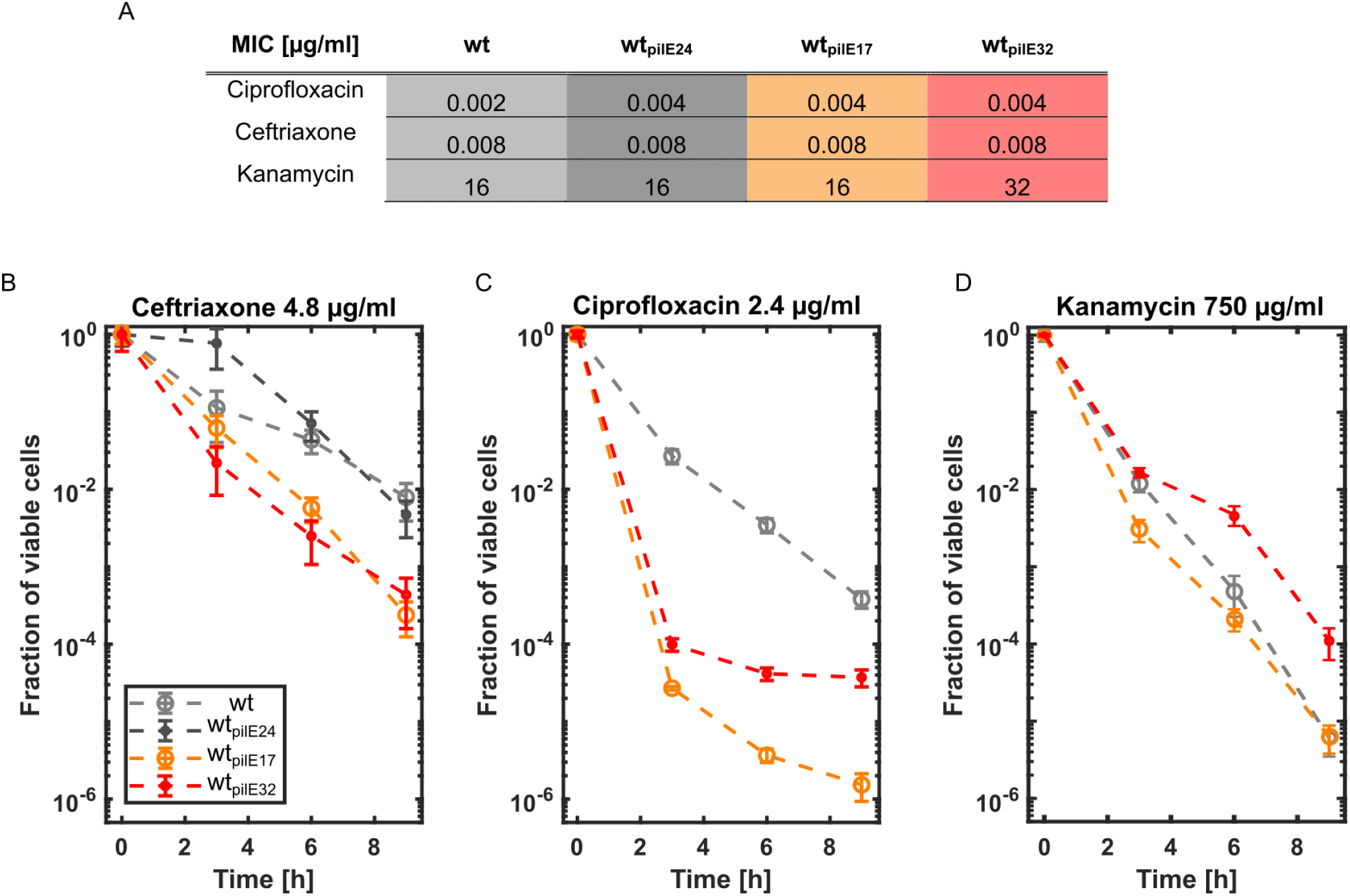
Effects of colony formation on antibiotic resistance and tolerance. A) MICs for all pilin variants and antibiotic treatments were determined as the modal value from three biological replicates. Fraction of viable cells as a function of time during treatment with B) ceftriaxone (4.8 µg/ml), C) ciprofloxacin (2.4 µg/ml), and D) kanamycin (750 µg/ml). N = 3. Error bars: standard error.

To characterize antibiotic tolerance, we examined the killing kinetics during treatment with lethal doses of antibiotics. Ceftriaxone is currently recommended for treatment of gonorrhea [41] and, therefore, we started by investigating its effects on survival during the planktonic and colony lifestyles. We let gonococci grow for 10 h in solution as described above and then added ceftriaxone at 4.8 µg/ml, which is 600x the MIC. We found that the fraction of viable colony forming cells (strains wt* and wt*_pilE24_*) was higher than the fraction of viable planktonic strains (wt*_pilE17_* and wt*_pilE32_*) (Fig. 5D), suggesting that colony formation enhances tolerance against ceftriaxone.

To verify that the antibiotic concentration is not a limiting factor for killing, we investigated the time-kill kinetics at three different ceftriaxone concentrations (2.4 µg/, 4.8 µg/ml and 9.6 µg/ml) and found that the colony forming strains are more tolerant than the planktonic strains independent of the ceftriaxone concentration (Fig. S7A-C). Concluding, tolerance is not due to the colony-forming cells having a lower antibiotic dose per cell, as their cell density is in fact lower after 10 h of growth than that of the planktonic cells, meaning their antibiotic dose per cell is even higher (Fig. 4A, B). This suggests that we might underestimate the protective effect of colony formation. Since the killing kinetics were qualitatively independent of the antibiotic concentration, we conclude that the antibiotic dose per cell plays a minor role in this assay. Next, we tested whether the growth phase affects the protective effect of colony formation (Fig. S8). We treated one of the planktonic strains, wt*_pilE17_*, and one of the aggregating strains, wt*, at 6 h of growth, i.e. in the middle of exponential growth with ceftriaxone at 600x MIC (4.8 µg/ml). We observed that the protective effect of aggregation, observed for wt*\**, was even stronger compared to treatment after 10 h of growth. We conclude that colony formation makes gonococci more tolerant to ceftriaxone treatment.

Finally, we addressed tolerance against bactericidal antibiotics with different cellular targets, in particular DNA gyrase/topoisomerase (ciprofloxacin), and the ribosome (kanamycin) at 10 h of growth (Fig. 5C, D) [42,43]. For these antibiotic treatments, we only investigated wt*, wt*_pilE17_* and wt*_pilE32_* because wt* and wt*_pilE24_* displayed similar phenotypes in our assay. We tested ciprofloxacin at 600x MIC (2.4 µg/ml) and kanamycin at 46x MIC (750 µg/ml) as the latter is not soluble at concentrations corresponding to 600x MIC. For ciprofloxacin (Fig. 5C), the difference between the aggregating and the planktonic strains is even more pronounced than for ceftriaxone, with colony-forming wt*\** exhibiting substantially higher viability than planktonic wt*_pilE17_* and wt*_pilE32_* cells. Notably, the planktonic cells showed a bi-phasic killing curve reminiscent of persister cells [32]. However, this effect was not observed with kanamycin treatment, with wt*_pilE32_* showing greater tolerance than wt* (Fig. 5D). Nonetheless, the observed differences in tolerance to ceftriaxone and ciprofloxacin show that antigenic variation impacts the survivability of *N. gonorrhoeae* by modulating T4P-mediated colony formation and growth

## Discussion

This study explored the effect of pilin antigenic variation on the biophysical characteristics of the Type 4 pili and its interplay with bacterial survival under antibiotic treatment. We reveal how antigenic variation governs the bacterial lifestyle, as hypermutable pilin variants possess distinct aggregating or planktonic phenotypes. Explicitly, *pilE* aggregative gonococcal variants exhibit a fitness advantage when treated with bactericidal antibiotics ceftriaxone and ciprofloxacin. Our results highlight the close relationship between aggregation and tolerance and demonstrate that antigenic variation plays an important role in bacterial survival and persistence that extends beyond escaping immune surveillance.

### Different pilin variants support different T4P functions

By investigating the pilin sequences of clinical isolates, we found PilE variants with distinct functionality. For example, PilE_17_ mediates twitching motility more efficiently and aggregation less efficiently than PilE_wt_. We demonstrate here that these differences are due to T4P stereochemistry and not to pilus density, in contrast to previous findings [7]. How might stereochemistry support different T4P-mediated functions? The undulating surface of the pilus may allow intimate interactions along the length of the pili via protruding knobs formed by the β-hairpins inserting into the holes between subunits. For strong pilus:pilus interactions these features would need to have stereochemical complementarity. From the structural models it is clear that even a single amino acid change in these knobs or holes can profoundly affect their shape and chemistry. Thus, a single amino acid change on one feature may require a compensatory change on the other for stereochemical complementarity to be maintained. PilE_wt_ and PilE_24_ are most similar in sequence. PilE_24_ is expected to have a glycan at Ser63 as does PilE_wt_ [44,24]. The glycan is positioned on the αβ-loop at the rim of the hole and impacts the opening size and chemistry. PilE_24_ differs from PilE_wt_ for a number of surface residues, both at the rim of the hole and on the knob, but since strain wt*_pilE24_* is a colony former like wt*, these differences are likely compensatory. Indeed, the knobs of PilE_wt_ and PilE_24_ appear to be somewhat negatively charged, which would complement their positively charged holes, but this could also be said of PilE_17_ and PilE_32_. Since the latter are not colony formers, it may be that while their knobs and holes are electrostatically complementary their shapes/structures are not. Kennouche et al. identified a positively charged patch in *N. meningitidis* PilE that is crucial for aggregation [16]. This region is centered on a conserved Lys140, which is located in the loop immediately following β-hairpin2. Though the *N. gonorrhoeae* pilins in this study do not possess a corresponding protruding lysine, this region is part of the β-hairpin knob and likely critical for pilus:pilus interactions. Our results suggest that small changes in the amino acid sequence of the major pilin subunit can have a large impact on pilus:pilus interactions and hence, *N. gonorrhoeae* behavior.

### The complex growth kinetics of gonococcal colonies

It has been reported previously that loss of T4P enhances the growth rate of *N. gonorrhoeae* [14]. This increase can be caused by the fact that pilin generation or pilus biogenesis consumes energy, reducing the growth rate [9] or by the fact that piliated gonococci form colonies in which central bacteria are growth-arrested [45]. In this study, all strains are comparable in their piliation levels, yet the colony formers wt and wt*_pilE24_* demonstrate very different growth behavior from the non-colony formers wt*_pilE17_* and wt*_pilE32_*. During the initial exponential growth phase, colony-forming cells exhibit a slower growth rate than planktonic cells, consistent with the reduced growth rate of bacteria residing at the center of colonies [45]. After ∼ 12 h, the colony-forming bacteria show a rapid increase in optical density, which is a good indicator for dissociation of *Neisseria* [49]. We interprete the rise in optical density as colony disassembly, releasing planktonic bacteria. Disassembly is most likely triggered by oxygen depletion as shown previously [46]. Oxygen concentration affects gonococcal behaviour in a complex way. For example, it governs membrane potential [47,48], T4P dynamics [49], and upregulation of the (truncated) denitrification pathway [50,51]. Since local oxygen gradients form within colonies [49], the oxygen availability is very different between colony-forming and planktonic cells. It is conceivable that the transient stationary phase of colony-formers indicates the period of time where the denitrification pathway is upregulated. Following this transient period of growth arrest, cells may resume growth employing denitrification. Understanding the growth kinetics is important for the interpretation of antibiotic tolerance discussed in the following paragraph.

### Pilin antigenic variants are tools for studying the mechanisms for colony-related tolerance

It is widely accepted that biofilm formation leads to higher tolerance against antibiotic treatment [51–53], but the mechanisms causing tolerance are poorly understood. *N. gonorrhoeae* form colonies in solution that have properties similar to bacteria biofilms, including a gradient of limited oxygen, of growth, and of tolerance agains antibiotics [45][47], and thus allow a systematic characterization of the effects of bacterial aggregation on tolerance. Here, we established stable *N. gonorrhoeae* strains expressing non-aggregative T4P that are otherwise fully functional, thus serving as ideal planktonic control strains.

We have addressed tolerance and resistance against three different antibiotic classes by comparing the killing kinetics of the colony-forming and planktonic control strains. The minimal inhibitory concentrations were minimally affected by colony formation, indicating that aggregation does not contribute to antibiotic resistance. In contrast, we show that T4P-mediated aggregation has a strong effect on tolerance. For ceftriaxone, we found an order of magnitude increase in the fraction of viable cells when bacteria formed colonies. This effect was robust with respect to the antibiotic concentration and the growth phase. Ceftriaxone is a β-lactam and for this class of antibiotics it has been shown for *E. coli* that the killing rate is inversely correlated with growth rate [52]. Similarly, growth-rate dependent killing rates were reported for *E. coli* treated with ciprofloxacin [53]. Bacteria at the center of gonococcal colonie are growth-arrested [45], and the fraction of dead cells in this location is lower under ceftriaxone treatment compared to the edges of the colonies [26], suggesting that local growth arrest enhances tolerance. Moreover, we propose that the formation of oxygen gradients within gonoccal colonies [47] can protect bacteria residing at the colony center where the oxygen concentration is lower. There is evidence that bactericidal antibiotics including ciprofloxacin and ceftriaxone kill bacteria (at least partially) by producing reactive oxygen species [54,55]. Reduced antibiotic penetration is suggested to enhance tolerance [56]. We can exclude this mechanism for our system, since we showed previously that antibiotic treatment causes swelling of the cell body and this effect is homogenous throughout the colonies [26].

Unexpectedly, colony formation did not protect gonococci from kanamycin treatment. For early-stage colonies, we found that cells at the center of the colonies are more tolerant to kanamycin than peripheral cells, most likely because cell at the center have lower electrical membrane potential [47]. On the hand, central cells grow more slowly and it has been shown that antibiotics that irreversibly bind the ribosome (like kanamycin) are more effective for slow-growing bacteria [57]. It is unclear, however, whether a similar argument holds for the killing rate at kanamycin concentrations exceeding the MIC. It is also interesting to note that strain wt*_pilE32_* is more tolerant to ciprofloxacin and kanamycin than strain wt*_pilE17_*, indicating that pilin antigenic variation affects tolerance by a mechanism independent of aggregation. It will be interesting to examine this additional effect in future studies. Taken together, the T4P is an important determinant of antibiotic tolerance as it governs aggregation and colony formation. Reduced growth rate and reduced oxygen concentration are likely involved in aggregation-associated tolerance against ceftriaxone and ciprofloxacin.

## Conclusion

Based on our results we propose that pilin antigenic variation has functions beyond its well-known role in escape from immune surveillance. Within a gonococcal population, antigenic variation rapidly generates a standing variation of different *pilE* sequences with phenotypes that support adhesion, aggregation, twitching motility, or DNA uptake. While the generation of variations is likely random, variants with different phenotypes are selected for during infection by the host environment. Here, we showed that these phenotypic changes impact antibiotic tolerance and, therefore, antigenic variation plays a key role in colony diversification, infection, and treatment of gonorrhea.

## Acknowledgements

We thank the CECAD Imaging Facility (and Felix Gaedke) for their support (in microscopy/technique / data analysis), and Rosalind Allen, Isabel Rathmann, Stephan Wimmi, and the Maier group for helpful discussions. This work has been supported by the Center for Molecular Medicine Cologne, the Deutsche Forschungsgemeinschaft through grant MA3898, and the the IHRS BioSoft.

## Supplementary Materials and Methods

### Growth conditions

We used the same growth conditions described in previous studies [26]. Gonococcal base agar was made from 10 g/l dehydrated agar (BD Biosciences, Bedford, MA), 5 g/l NaCl (Roth, Darmstadt, Germany), 4 g/l K_2_HPO_4_ (Roth), 1 g/l KH_2_PO_4_ (Roth), 15 g/l Proteose Peptone No. 3 (BD Biosciences), 0.5 g/l soluble starch (Sigma-Aldrich, St. Louis, MO), and supplemented with 1% IsoVitaleX (IVX): 1 g/l D-glucose (Roth), 0.1 g/l L-glutamine (Roth), 0.289 g/l L-cysteine-HCL x H_2_O (Roth), 1 mg/l thiamine pyrophosphate (Sigma-Aldrich), 0.2 mg/l Fe(NO_3_)_3_ (Sigma-Aldrich), 0.03 mg/l thiamine HCl (Roth), 0.13 mg/l 4-aminobenzoic acid (Sigma-Aldrich), 2.5 mg/l β-nicotinamide adenine dinucleotide (Roth), and 0.1 mg/l vitamin B12 (Sigma-Aldrich). GC medium is identical to the base agar composition but lacks agar and starch.

### Bacterial strains

Clinical isolates NG17, NG24 were cultured from urethral swabs in 2016, and NG32 in 2017, from male patients presenting with urethritis. After identification and susceptibility testing, they were stored in glycerol at -80°C. Isolates were recovered from the freezer by plating out on chocolate agar for the purpose of this study.

First, clinical isolates were transferred back into the piliated state as described in the following. We grew bacterial cells for 1 day in liquid medium (37 °C and 5% CO_2_) without shaking. The next day we transferred bacteria growing at the surface with a loop to fresh GC-media. This process was repeated until a pellicle at the surface was formed. Then we plated the pellicle on GC agar plates. Piliated phenotypes were chosen according to the colony morphology and testing for twitching motility.

To determine the sequence of the *pilE* variants we performed a PCR with primers sk5 and sk34 on the respective gDNA (isolated with the Blood and Tissue Kit, Qiagen). The PCR product was sequenced with primer pilE_IWupstream_ (Eurofins).

### Construction of *pilE* variant strains

In order to construct isogenic strains that differ solely in the *pilE* sequence, the *pilE* sequences of the clinical isolates were cloned into the ΔG4 background strain (Ng150, derivative of MS11, [58]) replacing the native *pilE* gene. To avoid further genetic modification the *pilE* genes were introduced by *ermC-rpsL_s_* based clean insertion as described [17].

The process for the construction was identical for strains wt_pilE17_, wt_pilE24_, and wt_pilE32_ except for the respective primers. First, the 5’ UTR region of *pilE* was amplified from gDNA of strain Ng150 using primers sk159 and sk160. Second, the *pilE* gene was amplified from gDNA of the different clinical isolates NG17, NG24, and NG32 with primers sk161+sk175, sk161+sk167 and sk161+sk162, respectively. Third, the 3′UTR of *pilE* including the *ermC-rpsL_s_* was amplified from gDNA of strain Ng225 using primers sk163+sk158. The three PCR products were fused and the final product was spot transformed into strain Ng150. After selection on erythromycin, insertions were controlled via screening PCR with primers sk32 and sk45. The respective strains were named *ΔG4 pilE*NG17/24/32 clean insertion step 1 (Ng239, Ng241, Ng229, respectively).

Next, the fusion construct for the counter selection was generated. To this end, the 5′ UTR region including the newly introduced *pilE* genes was amplified from gDNA of strains *ΔG4 pilE*NG17/32/24 clean insertion step 1 (Ng239, Ng229, Ng241) with primers sk159+sk176, sk159+ sk168, sk158+sk165, respectively. The 3′ UTR of *pilE* was amplified from gDNA of strain Ng150 with primers sk164+sk158. The two products were joined in a fusion PCR and transformed into the respective strain of the first step (Ng239, Ng241, Ng229). During selection on streptomycin the *ermC*-*rpsL*_s_ construct is spliced out and the mutants are isogenic to the parental strain (Ng150) except for the *pilE* sequences. Insertions were controlled via screening PCR with primers sk32 and sk45 and subsequently checked via sequencing with primer sk129 (Eurofins). The final strains are referred to as wt*_pilE24_* (Ng242), wt*_pilE17_* (Ng240), wt*_pilE32_* (Ng230).

### Twitching motility analysis

We let the strains grow for 12-16 h on GC agar plates. We picked a few colonies, resuspended them in liquid GC medium, transferred them to a BSA coated coverslip (1 mg/ml), and then sealed the sample with Valep (Vaseline, wool fat and paraplast in a ratio of 1:1:1). Subsequently, we recorded videos with a confocal Ti-E inverted microscope (Nikon) equipped with a thermobox at 37°C with a framerate of 10 Hz and 100 x magnification over 30 s. We stopped each measurement after 15 min to ensure constant experimental conditions.

Next, we tracked each bacterium over the whole measurement time. From each track, we calculated the MSD from the displacements 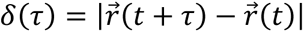 (MATLAB 2021). By averaging over all tracks of each strain, we determined the average MSD curve. As the bacteria follow a correlated random walk, we fitted 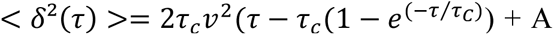 for the first five seconds to the data to determine the correlation time τ_c_ and velocity *v*. The variable A accounts for the tracking error. During the fitting procedure, the data was weighted by its standard deviation (Fig. S5).

### Transmission electron microscopy and determination of T4P number

Bacteria grown overnight on GC agar plates were resuspended in liquid medium and adjusted to an optical density (OD_600_) of 0.1. For sample preparation, 10 µl of the bacterial solution was transferred on a 100 mesh formvar coated copper grid (Science Services) and incubated for 20 min at room temperature. For fixation, the grid was put upside down in a drop of 2% formaldehyde (Science Services) and incubated for 5 min, followed by five times washing in PBS. Next, the mesh was placed on a drop of 1% glutaraldehyde (Sigma) for 5 min and washed eight times in Milli-Q water. The samples were blotted on filter paper. For negative staining, the cells were again incubated with 10 µl of uranyl acetate for 4 min and the spill-over was removed by filter paper. Then, they were imaged in a transmission electron microscope (JEM-2100Plus (JEOL)) in the imaging facility of the CECAD, Cologne. All images were taken at 6000 x magnification at 40 µm under the default focus at room temperature.

The numbers of T4P per cell were counted manually. We note that T4P form bundles in which single T4P cannot be resolved properly and T4P sometimes overlap. Therefore, the number of T4P per cell is considered an estimate. Compared to our previous study, the T4P density was considerably higher [20]. Here, we used a different growth medium that supports gonococcal growth better and explains the higher level of piliation.

### Generation of PilE and pilus models

AlphaFold2 [39] was used to generate models of PilE_wt_ and the PilE varients. Models were superimposed upon Chain A of the *N. gonorrhoeae* T4P cryoEM reconstruction [38] in Chimera (Petterson et al., PMID 15264254) and residues 1-48 of the model were replaced with that of Chain A to replace the continuous α1-helix of the AF2 models with the melted helix seen in the filament. The helical symmetry parameters of *N. gonorrhoeae* T4P (10.1 Å rise, 100.8° rotation) were imposed on the PilE models to generate 18-mer filament models. Electrostatic potential was generated using the APBS tool of PyMol [59].

### Confocal microscopy

The local order and within-colony motility of the gonococcal colonies were determined by following a previously developed protocol [26]. Briefly, we resuspended bacteria from overnight GC agar plates in GC liquid media to an optical density OD_600_ 0.1. Subsequently the bacterial solution was incubated for 45 min in a shaking incubator to let the cells aggregate. Next, we inoculated 300 µl of the suspension into a Poly-L-lysine coated (Sigma, stock solution 50 μg/ml) Ibidi 8-well plate, stained the bacteria with 2 µmol/l Syto-9 (Thermo Fisher Scientific) and incubated them for another ten minutes to stain all bacteria homogenously. All images were acquired using an inverted microscope (Ti-E, Nikon) equipped with a thermo box (37°C) and a spinning disk unit (CSU-1, Yokogawa) with 100x magnification, 1.49 NA, oil immersion objective lens. The excitation wavelength was 488 nm with 0.5% laser power. We imaged the samples for 30 minutes, as we observed that the colonies showed disintegration after this time.

To analyse the within-colony motility, we acquired images in a 2D-focal plane with a framerate of 10 Hz. Three-dimensional z-stacks with a plane-to-plane distance of 0.2 µm and an overall height of 10 µm were acquired to investigate the local order of colonies.

### Single-cell tracking in three dimensions and analysis of the local order of microcolonies

The spatial analysis in three dimensions was performed with a program already implemented for previous publications [21,26]. In short, stacks of confocal images were filtered with a spatial bandpass filter. The resulting image showed local maxima which were identified as bacterial coordinates with pixel accuracy. Next, spherical masks around each local maximum were created and the subpixel positions were calculated from the centroids. From the bacterial coordinates we determined the radial distribution function *g(r)* which is defined as *N/V g(r)r^2^dr*, the probability to find the next bacterium in a distance *r* within a shell *dr*. *r* is the center-to-center distance and *N/V* displays the density of the gonococcal colony. To determine *g(r)*, we calculated the distances between all pairs of bacteria and sorted them according to the distances with bins of width *dr*, resulting in a distribution of distances. By further processing this distribution via smoothing and average filtering, we generate a general distribution of particles with the same particle density but random distances, *N(r)*. The radial distribution function was determined via *g(r) = n(r)/N(r)*. From the radial distribution function, it was shown that the course of the radial distribution reveal information of the colloidal system with Lennard-Jones-like interaction, as described previously [21].

### Measurement of within-colony motility

We followed a protocol which was described in [26]. We first tracked the cells in the focal plane in Image J Trackmate [40], following the procedure described in [21] Then, for each track longer than 3 s the MSD was calculated. Next, we determined the diffusion constants *D* from the resulting MSDs with *D* = *MSD*(*t* = 1 *s*)/(4 · 1*s*) and corrected the static errors via the velocity auto-correlation function as described in [60].

### Dual laser tweezers experiments

The interaction forces of *pilE* variants were determined via a dual laser trap. The experimental setup and analysis is already published [21]. In short, we resuspended a few bacterial colonies from overnight GC agar plates in liquid GC medium. We added 1:1000 ascorbic acid (500 mM). Next, we inoculated the bacteria on a BSA coated cover slip (1 mg/ml) and sealed the slide with VALEP (Vaseline, wool fat and paraplast in a ratio of 1:1:1. The major building blocks of the laser tweezers setup consist of a microscope equipped with a thermo-box at 33°C, an IR-laser (1064 nm) and an acousto-optical deflector which creates two time-shared optical potentials. The trap distance was set to 2.64 µm. We acquired videos of interacting bacteria with a framerate of 50 Hz. After 15 min, we stopped the measurement as we observed decreased activity of the gonococci.

We detected the displacements *d* of the bacteria from the equilibrium positions via a Hough transformation algorithm. From the displacement tracks we determined the forces (*F ∼ d*) and identified the interaction states, as described earlier in [21]. The potential of each trap was approximated to be harmonic for forces up to 80 pN.

At 100% laser intensity, the traps showed a trap stiffness of *k*_100%_= 0.1 pN/nm whereby the laser intensity *I* is proportional to the stiffness of the trap *k* ∼ *I*. We note that it was not possible to conduct the experiments with the same laser intensities for all variants, since deflections of the wt*_pilE32_* and wt*_pilE17_* strains were infrequent. This indicated that these two strains have lower interaction forces as they could not overcome the trapping potential of the traps at 100% laser power. Therefore, we adjusted the intensities of the laser. The measurements for the wt* and wt*_pilE24_* could be performed at 100% laser power while for the other variants, the laser intensity needed to be decreased to 10% for strain wt*_pilE32_* and to 5% for strain wt*_pilE17_*. Under these conditions, the probability of pilus:pilus binding (Fig. S2) was high enough for characterizing T4P mediated attractive forces.

### Bacterial growth curves

Bacterial growth and aggregation were monitored by measuring the OD600 with an Infinite M200 plate reader. After 12–14 h on GC-agar plates, bacteria were resuspended in liquid GC medium and adjusted to an optical density OD_600_ of 0.1. For each time point and each condition, a 48-well plate (Greiner), containing 1 ml liquid GC media, was inoculated with 10 µl of the bacterial suspension. We incubated the bacteria at 37°C, 5% CO_2_ with a shaking period of 2 min per OD cycle, and measured the OD every 10 min.

To determine the number of colony forming units (CFU) during 19 h of growth, we performed the same protocol as described above and additionaly transferred a whole well to a 1.5 ml reaction tube every 1-3 h. Next, we harvested the bacteria by centrifugation (5000 g, 3 min) and resuspended them in 500 µl GC media. Subsequently, we added 500 µl of MQ-water to initiate the disassembly of gonococcal aggregates and vortexed the suspension for 2 min. We performed 1:10 dilution series and plated 50 µl of different dilutions on non-selective GC agarplates. After 48 h of growth (37°C with 5% CO_2_) we counted the CFUs.

### MIC determination

The minimal inhibitory concentration of the different antibiotics (ceftriaxone, ciprofloxacin, kanamycin) was determined for each strain. 1 ml cultures supplemented with increasing antibiotic concentrations were inoculated with approximately 5 · 10^5^cells of the following strains, wt* (Ng150), wt*_pilE24_* (Ng242), wt*_pilE17_* (Ng240), and wt*_pilE32_* (Ng230). Bacteria were grown in an Infinite M200 plate reader at 37°C, 5% CO_2_ with a shaking period of 2 min per OD cycle. The lowest concentration of an antibiotic without detectable growth (OD_600nm_ ≤ 0.1) after 24 h was determined as the MIC of the respective antibiotic.

### Bacterial survival assay

To investigate how antigenic variation impacts bacterial survival under antibiotic treatments we developed a survival assay. Isolates were initially grown as described for the bacterial growth curves. Following resupension in GC media, we let the bacteria grow for 10 h in an Infinite M200 plate reader at 37°C, 5% CO_2_ with a shaking period of 2 min per OD cycle. If other pre-growth durations were used, we indicated this in the figure and description for the specific experiments. OD was measured every 10 min. Next, we added antibiotics to each well except the control wells, and the plate was further incubated. The final concentrations of ceftriaxone were 2.4/4.8/9.6 µg/ml corresponding to the 300x MIC/600x MIC/1200x MIC, respectively. For ciprofloxacin treatment, we added antibiotics to a final concentration of 2.4 µg/ml, and for kanamycin, the concentration was 240 µg/ml, corresponding to 600x MIC and 46x MIC, respectively. The solubility of kanamycin was too low to increase the antibiotic concentration. To determine the number of viable bacteria, cells were plated at 0 h, 3 h, 6 h, and 9 h after antibiotic treatment as described before (Methods: Bacterial growth curves).

## Supplementary Figures

**Fig. S1.**
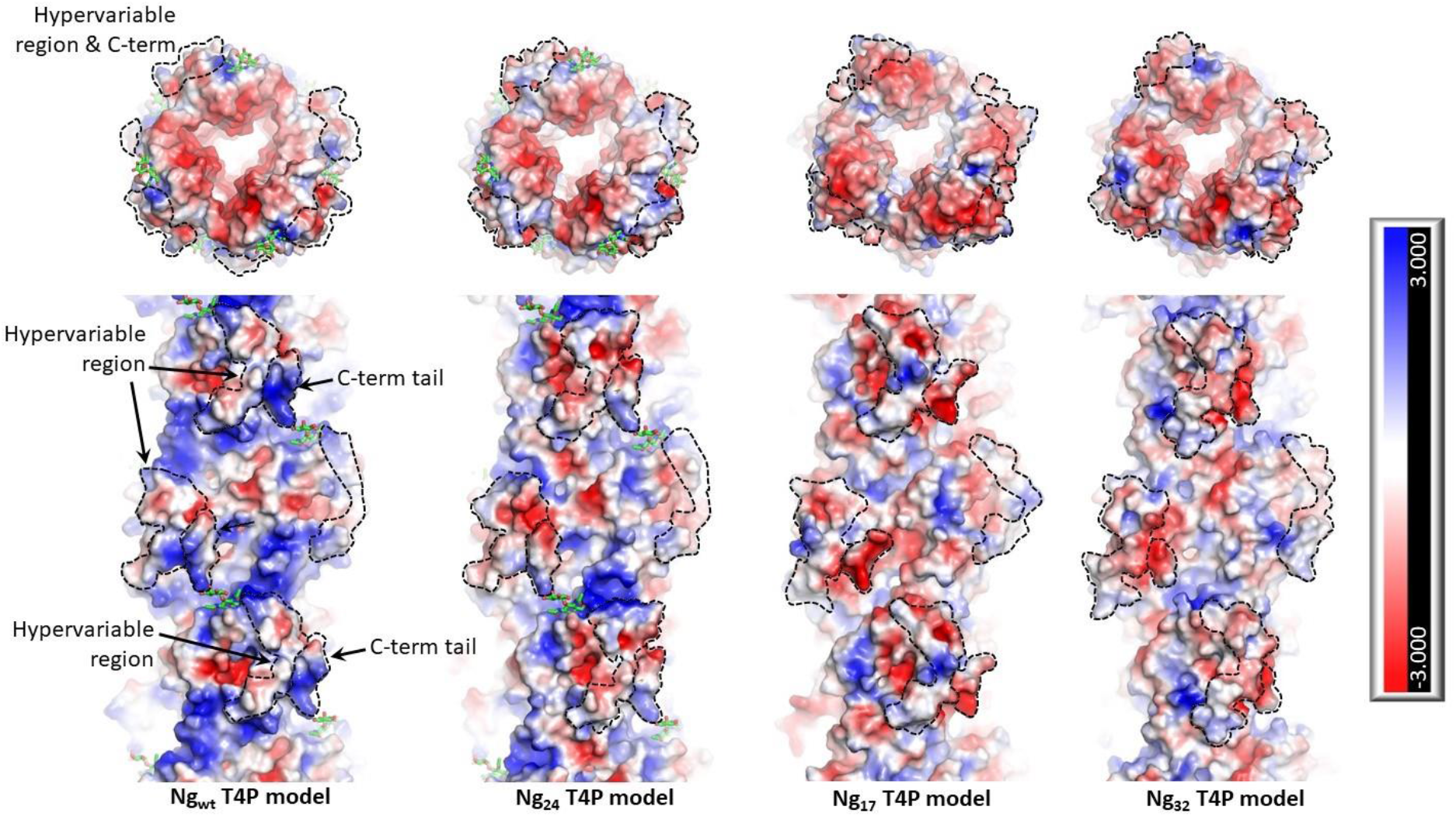
Model of the charge densities. The charge density was simulated via PyMol and the APBS tool [59][61]. Models of wt* and variant pilus filaments are shown from the top and side in surface representation with electrostatic surface potential. Blue: positive charge, red: negative charge.

**Fig. S2.**
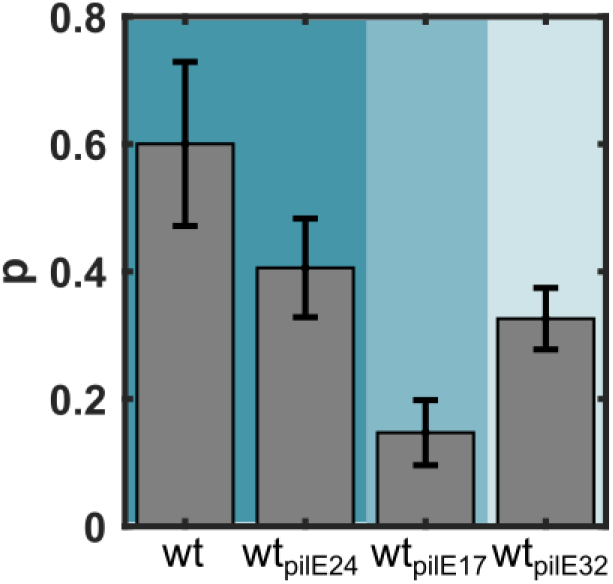
Fraction of successful attempts in dual trap assay. We counted the fraction of interacting bacteria pairs because not every pair of bacteria showed interaction. This fraction strongly depends on the trap stiffness which was set to *k* = 0.1 pN/nm for wt*\** and wt*_pilE24_* (dark blue). Since interactions were nearly undetectable at *k* = 0.1 pN/nm for strains wt*_pilE32_* and wt*_pilE17_*, the stiffnesses were reduced to *k* = 0.005 pN/nm (light blue) and k = 0.001 pN/nm (blue), respectively. Number of trapped bacteria pairs: N_wt_ = 73, N_pilE24_ = 167, N_pilE17_ = 170, N_pilE32_ = 168. Error bars: standard error over the measurement days.

**Fig. S3.**
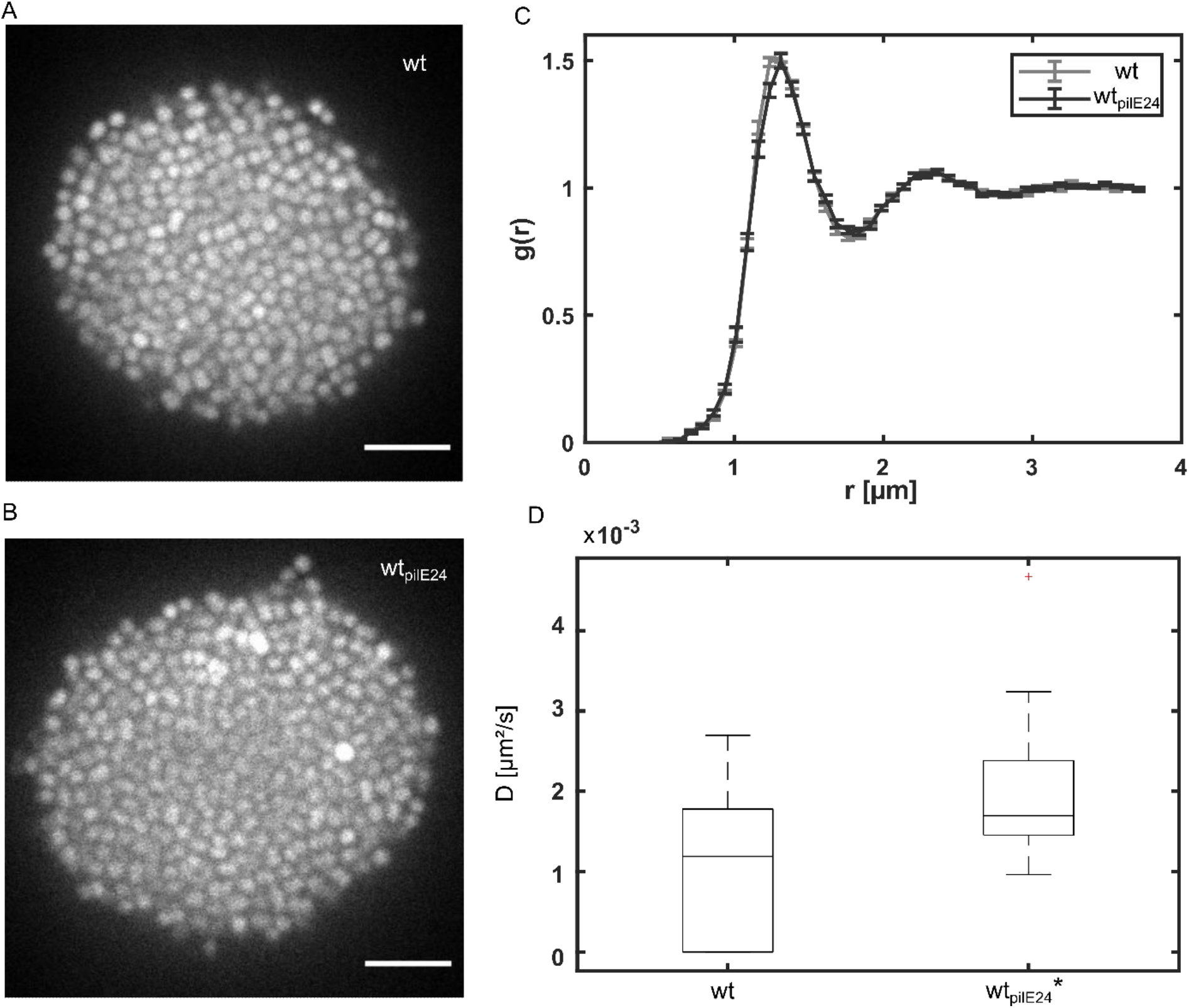
Colony structures of *wt\** and *wt_pilE24_* variants. A) Typical confocal planes through the centres of gonococcal colonies for strain *wt\** (A) and *wt_pilE24_* (B) stained with 2 mM Syto-9. C) Radial distribution functions of the aggregating strains. Grey: *wt\** and dark grey: *wt_pilE24_*. N=21-27. D) Diffusion constants D of single cells 8 µm inside the colonies for *wt\** and *wt_pilE24_*. KS-test: p = 3.1×10^-5^. Box plot: median, 25/75 percentile, whiskers: interquartile range times 1.5. N = 18-21.

**Fig. S4.**
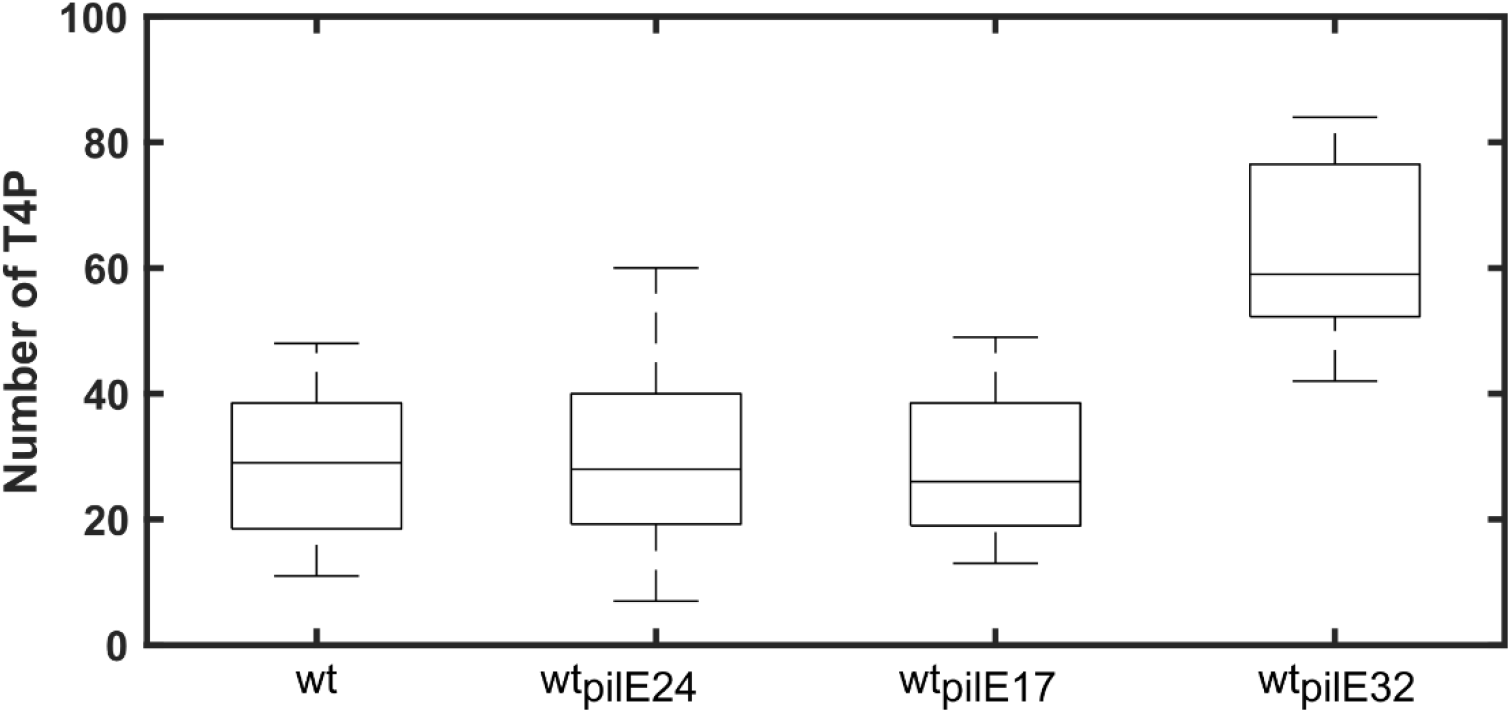
Number of T4P per cell for strains wt* (Ng150), wt*_pilE24_* (Ng242), wt*_pilE17_* (Ng240), wt*_pilE32_* (Ng230). N = 12 cells were investigated for each condition. Box plot: median, 25/75 percentile, whiskers: interquartile.

**Fig. S5.**
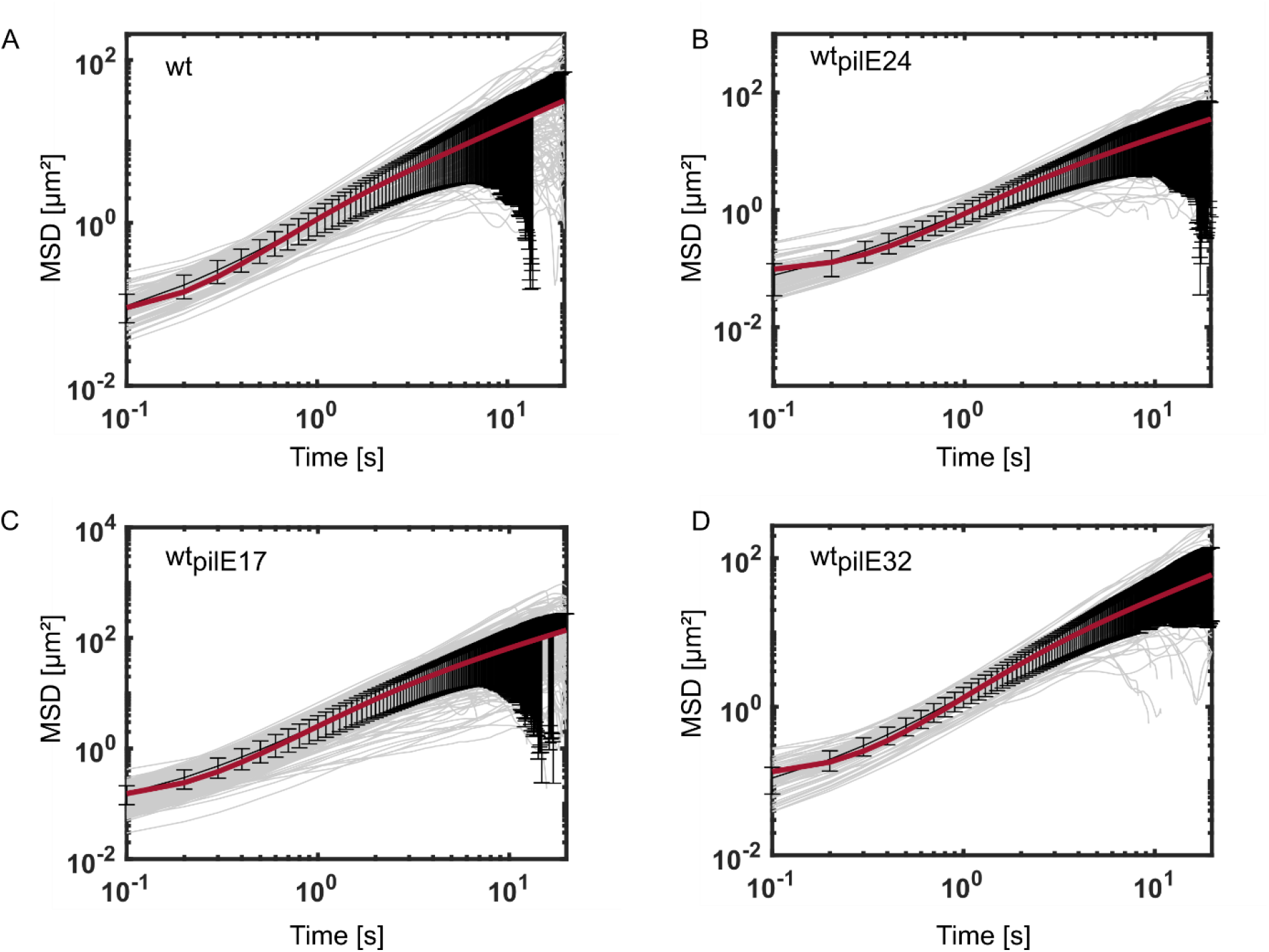
Mean squared displacement (MSD) for all tracks of single cells on a BSA-coated cover slide for strains wt* (Ng150), wt_pilE24_ (Ng242), wt_pilE17_ (Ng240), wt_pilE32_ (Ng230). The MSD was fitted for the time interval of the first 5 s. Grey: trajectories of individual cells, black: mean trajectories, red line: fit to the correlated random walk model, error bars: standard error of the mean. N = 88-268 trajectories per strain.

**Fig. S6.**
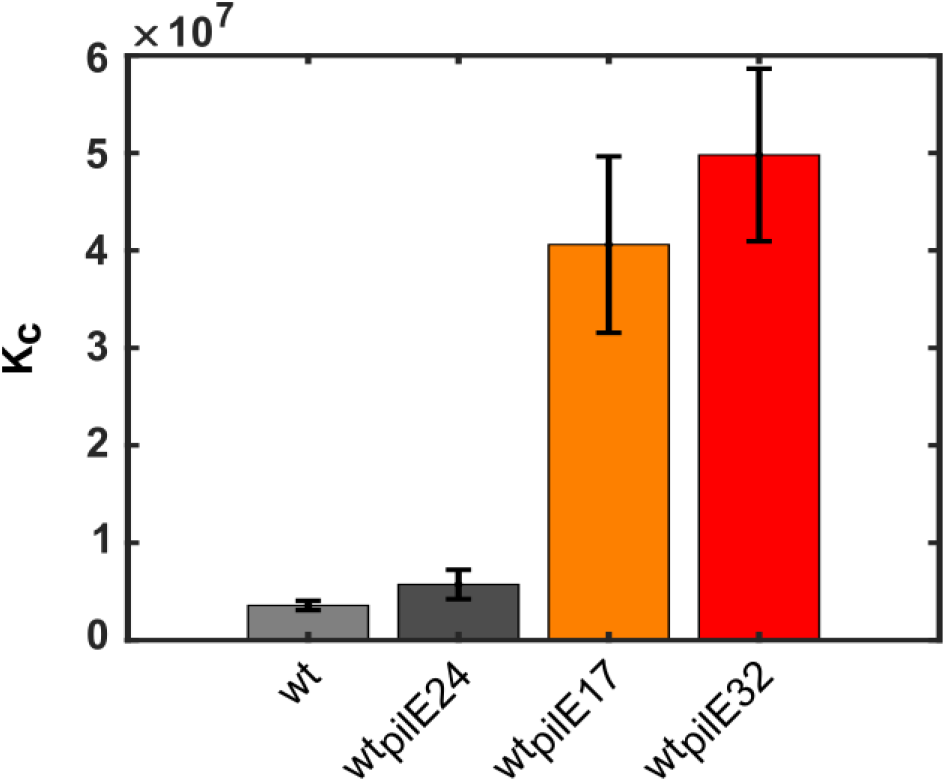
Carrying capacity is determined as the maximal number of cells of the first plateau of growth curve (Fig. 4). Error bar: standard error.

**Fig. S7.**
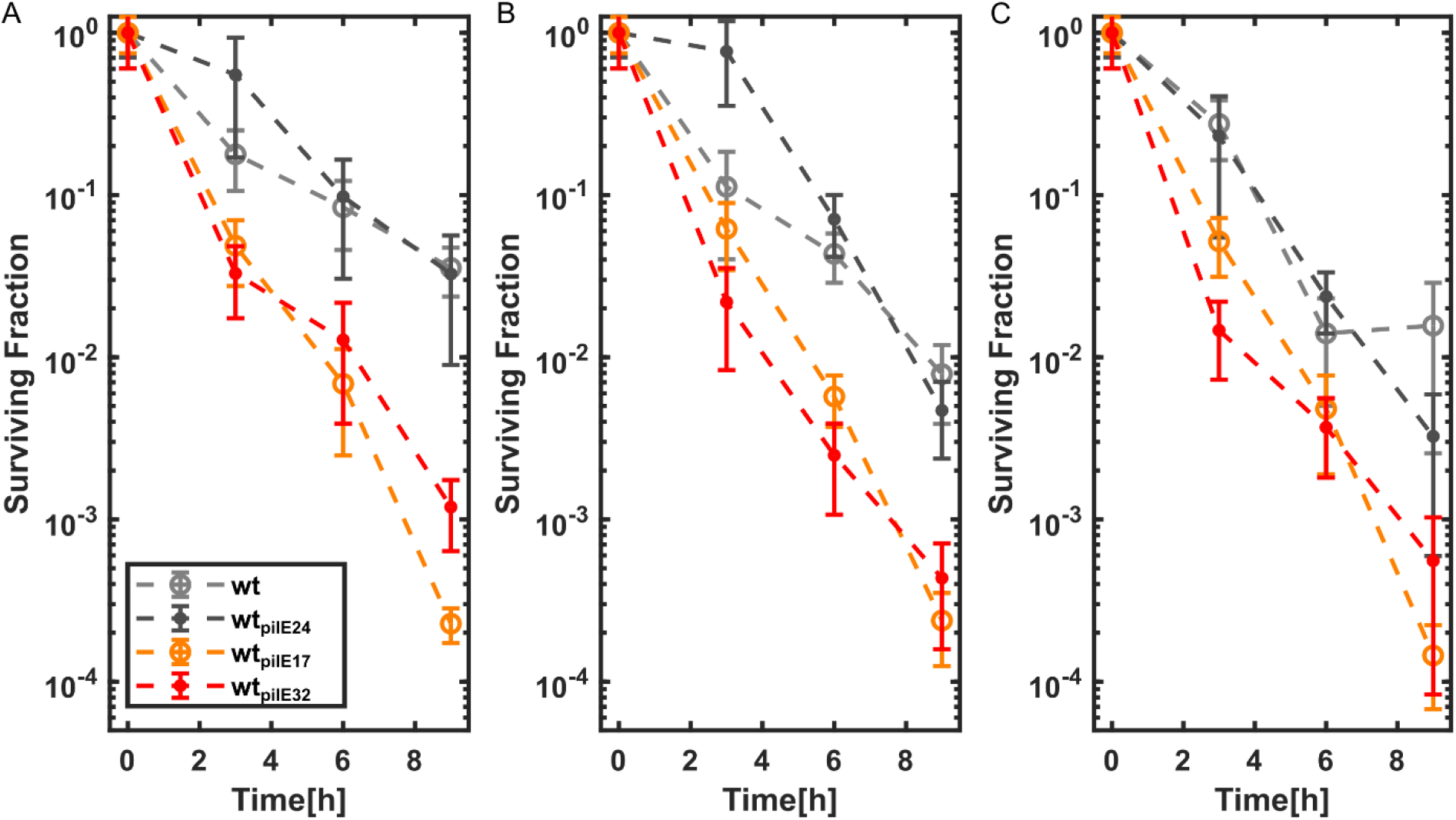
Survival assay with different concentrations of ceftriaxone after 10 h of growth. Killing kinetics of all variants for A) 300x MIC, B) 600x MIC, and C) 1200x MIC. Error bars: standard errors over 3 independent experiments.

**Fig. S8.**
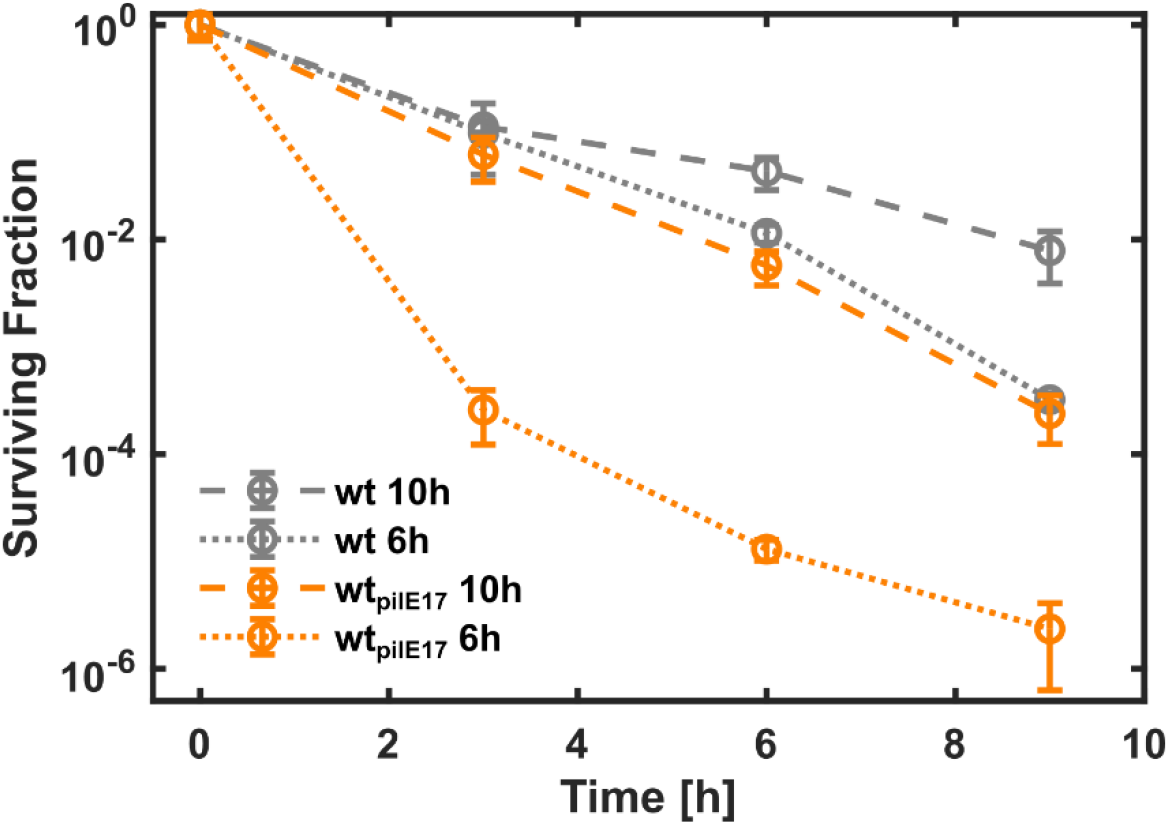
Killing kinetics in different growth phases. A) Killing kinetics of *wt\** and *wt_pilE17_* which grew either 6 h or 10 h before the start of the treatment with ceftriaxone at 600x MIC. Error bars: standard errors over 3 independent experiments.

## Supplementary Tables

**Table S1.** Strains used in this study.

| strain | relevant genotype | reference | strain data base entry |
| --- | --- | --- | --- |
| *N. gonorrhoeae* NG32 | - | clinical isolate | Ngc001 |
| *N. gonorrhoeae* NG17 | - | clinical isolate | Ngc004 |
| *N. gonorrhoeae* NG24 | - | clinical isolate | Ngc007 |
| $\Delta G4$ (wt*) | *G4::apraR* | [62] | Ng150 |
| *T126C* step1 | *G4::apraR, pilE::pilET126C ermC rpsLs* | [17] | Ng225 |
| *wt<sub>pilE32</sub>* step1 | *G4::apraR, pilE::pilEclinicalisolateNG32 ermC rpsLs* | this study | Ng229 |
| *wt<sub>pilE32</sub>* | *G4::apraR, pilEclinicalisolateNG32* | this study | Ng230 |
| *wt<sub>pilE17</sub>* step1 | *G4::apraR, pilE::pilEclinicalisolateNG17 ermC rpsLs* | this study | Ng239 |
| *wt<sub>pilE17</sub>* | *G4::apraR, pilEclinicalisolateNG17* | this study | Ng240 |
| *wt<sub>pilE24</sub>* step1 | *G4::apraR, pilE::pilEclinicalisolateNG24 ermC rpsLs* | this study | Ng241 |
| *wt<sub>pilE24</sub>* | *G4::apraR, pilEclinicalisolateNG24* | this study | Ng242 |

**Table S2.** Amino acid sequence identities of complete and partial regions of *pilE* compared to the MS11 *pilE* amino acid sequence according to Fig. 1. The amino acid sequences were compared using BLAST [63].

| Region | PilE variant | Query Cover [%] | Identity [%] | e-value | length |
| --- | --- | --- | --- | --- | --- |
| complete | NG17 | 98.0 | 80.6 | 4e-85 | 167 |
|  | NG32 | 96.0 | 86.6 | 5e-95 | 165 |
|  | NG24 | 99.0 | 90.0 | 3e-104 | 171 |
| without conserved | NG17 | 94.0 | 73.6 | 5e-48 | 115 |
|  | NG32 | 94.0 | 80.4 | 3e-57 | 113 |
|  | NG24 | 99.0 | 85.6 | 2e-66 | 119 |
| Semi-variable | NG17 | 100.0 | 76.1 | 1e-43 | 67 |
|  | NG32 | 100.0 | 80.6 | 2e-37 | 66 |
|  | NG24 | 100.0 | 89.6 | 2e-36 | 67 |
| cys1 | NG17 | 100.0 | 84.6 | 3e-13 | 13 |
|  | NG32 | 100.0 | 100.0 | 2e-15 | 13 |
|  | NG24 | 100.0 | 100.0 | 2e-15 | 13 |
| Hypervariable loop | NG17 | 94.0 | 57.9 | 5e-06 | 20 |
|  | NG32 | 100.0 | 59.1 | 1e-09 | 22 |
|  | NG24 | 100.0 | 56.5 | 2e-08 | 23 |
| cys2 | NG17 | 100.0 | 100.0 | 5e-12 | 10 |
|  | NG32 | 100.0 | 100.0 | 5e-12 | 10 |
|  | NG24 | 100.0 | 100.0 | 5e-12 | 10 |
| Hypervariable tail | NG17 | - | - | - | 5 |
|  | NG32 | - | - | - | 2 |
|  | NG24 | 83.0 | 100.0 | 5e-05 | 6 |

